# More than 16,000 effectors in the *Legionella* genus genome provide multiple, independent combinations for replication in human cells

**DOI:** 10.1101/350777

**Authors:** Laura Gomez-Valero, Christophe Rusniok, Danielle Carson, Sonia Mondino, Ana Elena Pérez-Cobas, Monica Rolando, Shivani Pasricha, Sandra Reuter, Jasmin Demirtas, Johannes Crumbach, Stephane Descorps-Declere, Elizabeth L. Hartland, Sophie Jarraud, Gordon Dougan, Gunnar N. Schroeder, Gad Frankel, Carmen Buchrieser

## Abstract

**Significance:** *Legionella pneumophila* is a bacterial pathogen causing outbreaks of a lethal pneumonia. The genus *Legionella* comprises 65 species for which aquatic amoebae are the natural reservoirs. Using functional and comparative genomics to deconstruct the entire bacterial genus we reveal the surprising parallel evolutionary trajectories that have led to the emergence of human pathogenic *Legionella.* An unexpectedly large and unique repository of secreted proteins (>16,000) containing eukaryotic-like proteins acquired from all domains of life (plant, animal, fungal, archaea) is contrasting with a highly conserved type 4 secretion system. This study reveals an unprecedented environmental reservoir of bacterial virulence factors, and provides a new understanding of how reshuffling and gene-acquisition from environmental eukaryotic hosts, may allow for the emergence of human pathogens.

**Abstract:** The bacterial genus *Legionella* comprises 65 species among, which *Legionella pneumophila* is a human pathogen causing severe pneumonia. To understand the evolution of an environmental to an accidental human pathogen, we have functionally analyzed 80 *Legionella* genomes spanning 58 species. Uniquely, an immense repository of 16,000 secreted proteins encoding 137 different eukaryotic-like domains and more than 200 eukaryotic-like proteins is paired with a highly conserved T4SS. Specifically, we show that eukaryotic Rho and Rab GTPase domains are found nearly exclusively in eukaryotes and *Legionella* species. Translocation assays for selected Rab-GTPase proteins revealed that they are indeed T4SS secreted substrates. Furthermore, F/U-box and SET domains were present in >70% of all species suggesting that manipulation of host signal transduction, protein turnover and chromatin modification pathways, respectively are fundamental intracellular replication strategies for *Legionellae*. In contrast, the Sec-7 domain was restricted to *L. pneumophila* and seven other species, indicating effector repertoire tailoring within different amoebae. Functional screening of 47 species revealed 60% were competent for intracellular replication in THP-1 cells, but interestingly this phenotype was associated with diverse effector assemblages. These data, combined with evolutionary analysis indicate that the capacity to infect eukaryotic cells has been acquired independently many times within the genus and that a highly conserved yet versatile T4SS secretes an exceptional number of different proteins shaped by inter-domain gene transfer. Furthermore we revealed the surprising extent to which legionellae have co-opted genes and thus cellular functions from their eukaryotic hosts and provides a new understanding of how dynamic reshuffling and gene-acquisition has led to the emergence of major human pathogens.

## Introduction

Legionnaires’ disease or legionellosis is an atypical pneumonia caused by bacteria of the genus *Legionella.* Shortly after the discovery of *L. pneumophila* (1) it was reported that this bacterium is pathogenic for freshwater and soil amoebae of the genera *Acanthamoeba* and *Naegleria* (2). This finding led to a new perception in microbiology, whereby bacteria that parasitize protozoa can utilize similar processes to infect human cells. Sequencing and analyses of the *L. pneumophila* genome substantiated this idea, when it revealed the presence of a large number and variety of eukaryotic-like domains within the predicted proteome (3). Many of these proteins, termed effector proteins, were shown to be secreted into the host cell where they facilitate *Legionella* intracellular replication within a specialized compartment termed the *Legionella* containing vacuole (LCV) (3, 4). Overall, the type IV secretion system (T4SS), Dot/Icm, secretes more than 300 different effector proteins into the host cell and is indispensable for virulence of *L. pneumophila* (5–8). The presence of the Dot/Icm T4SS in other *L. pneumophila* strains and in selected *Legionella* species had also been reported (9–12) but recent genome scale studies of *Legionella* (13–15) indicated that the T4SS system is present in every *Legionella* strain analyzed.

Despite high conservation of the Dot/Icm system among different *Legionella* species, effector repertoires appear to vary greatly. An analysis of putative T4SS effectors of *L. longbeachae*, the second most frequent cause of Legionnaires’ disease, revealed that only about 50% of the virulence factors described in *L. pneumophila* were also present in the genome of *L. longbeachae* (16). Recently, Burstein *et al.* (14) analyzed 38 *Legionella* species using a machine learning approach to predict T4SS effectors and Joseph *et al.* (15) examined *Legionella* genome dynamics, both concluding that DNA interchange between different species is rare. However, still little is known about the potential of the different species to cause human disease and about the impact and the specific characteristics of the T4SS effectors on the evolution of new human pathogens within this environmental bacterial genus.

Here we present a comprehensive analysis of the *Legionella* genus genome, covering 80 *Legionella* strains belonging to 58 *Legionella* species and subspecies. We establish a pan-genus pool of putative T4SS effectors and show that this comprises over 16,000 proteins and identify more than 200 new eukaryotic-like proteins and 137 eukaryotic domains, including a unique class of putative bacterial Rab GTPases. We confirmed experimentally that a subset of these proteins translocate into the host cell upon infection. We conclude that the T4SS is highly conserved at the sequence level, but the effector proteins secreted are highly diverse.

## Results and discussion

### The *Legionella* genus genome is dynamic and characterized by frequent genetic exchange

We sequenced 58 *Legionella* species of which 16 were newly seqeunced, and analyzed them in combination with all publicly available genomes (80 genomes in total) (**SI Appendix, Table S1**). The *Legionella* genomes were extremely diverse, as the genome size varied from 2.37Mb (*L. adelaidensis*) to 4.88Mb (*L. santicrucis*), the GC content from 34.82% (*L. busanensis*) to 50.93% (*L. geestiana*) and the number of clusters of orthologous genes as defined with OrthoMCL was 17,992 of which 5,832 (32%) were strain specific (singletons) (Fig. 1A). Only 1,008 genes (6%) constituted the core genome (Fig. 1B), compared to an earlier analysis of 38 *Legionella* species, which found 16,416 clusters of orthologous and 1,054 core genes (14). The addition of 40 new genomes comprising 16 newly sequenced *Legionella* species in our study increased the number of orthologous gene clusters by over 1,576 and decreased the core genome by 46 genes, underlining the high diversity of the *Legionella* genus. This difference suggested that the *Legionella* genus pan-genome is far from fully described and that sequencing of additional *Legionella* species will increase the genus gene repertoire significantly. This was supported by the rarefaction curve that does not reach a plateau (Fig. 1C).

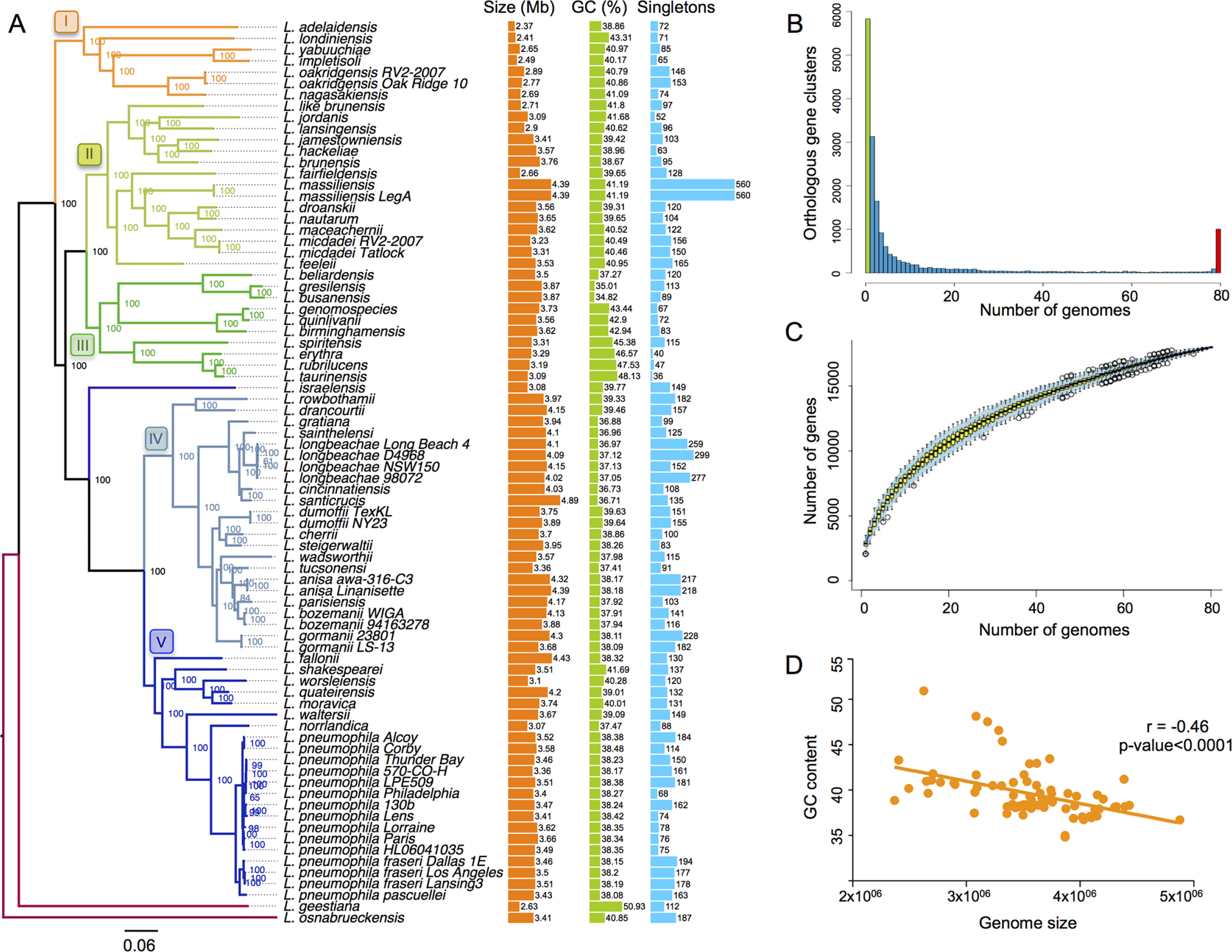

The highly dynamic nature of these genomes is also seen in the analysis of the strain specific genes and the accessory genome as it highlights the presence of several mobile genetic elements; often associated with genes encoding for transfer regions/conjugative elements such as the type IVA secretion systems (T4ASS). These T4ASSs (classified as T4SSF, G, I and T (17) are present in each strain to varying degrees indicating that they circulate among the different *Legionella* strains (**SI Appendix, Table S2**) and therefore drive genome dynamics and diversification. It has been suggested that the incorporation of foreign DNA via horizontal gene transfer (HGT) is responsible for an increase in the AT content and the increase in genome size (18). Indeed, we found a negative correlation between the genome size and the GC content for the *Legionella* genomes, which also suggests frequent HGT (Fig. 1D) (19). Despite the importance of flagella for transmission to new hosts as shown for *L. pneumophila*, flagella encoding genes were not conserved in all species, but showed a patchy distribution, as 23 of the 80 strains analyzed lacked flagella genes (**SI Appendix, Fig. S1**). The analyses showed that the *Legionella* genus genome is highly diverse, dynamic and shaped by HGT.

### The genus *Legionella* encodes proteins with 137 different eukaryotic domains

Interpro scan analysis of all 58 *Legionella* species revealed the presence of 137 different eukaryotic motifs/domains in the genus *Legionella* (**SI Appendix, Table S3**) according to the definition that an eukaryotic domain is one that is found in >75% of eukaryotic genomes and <25% in prokaryotic genomes. Interestingly, *L. santicrucis and L. massiliensis* encoded 41 and 39 ankyrin domains, respectively (Fig. 2). Ankyrin motifs were found frequently associated with other eukaryotic motifs and thus constituted modular proteins associated with eukaryotic F-box, U-box, Rab or SET domains. Notably, F-box and U-box domains were present in more than two thirds of the species analyzed (Fig. 2) suggesting manipulation of the host ubiquitin system is a fundamental virulence strategy of *Legionella* species. Generally, the genomes contained one to three F-box containing proteins with the exception of *L. nautarum* and *L. dronzanskii,* which contained 18 and 10, respectively. The SET domain containing protein RomA of *L. pneumophila* that induces a unique host chromatin modification (20) is present in 46 of the 58 *Legionella* species suggesting the ability of many *Legionella* species to manipulate host chromatin (Fig. 2). Interestingly, the Sec-7 domain present in the effector RalF, a bacterial ARF guanine exchange factor and the first described Dot/Icm effector of *L. pneumophila* (21) was present in only eight (*L. pneumophila, L. longbeachae, L. feelei, L. sainthelensis, L. santicrucis, L. shakespeari, L. quateirensis L. moravica*) of the 58 *Legionella* species analyzed, suggesting that, different effectors may compensate for RalF activity or that LCV biogenesis varies among different species (Fig. 2).

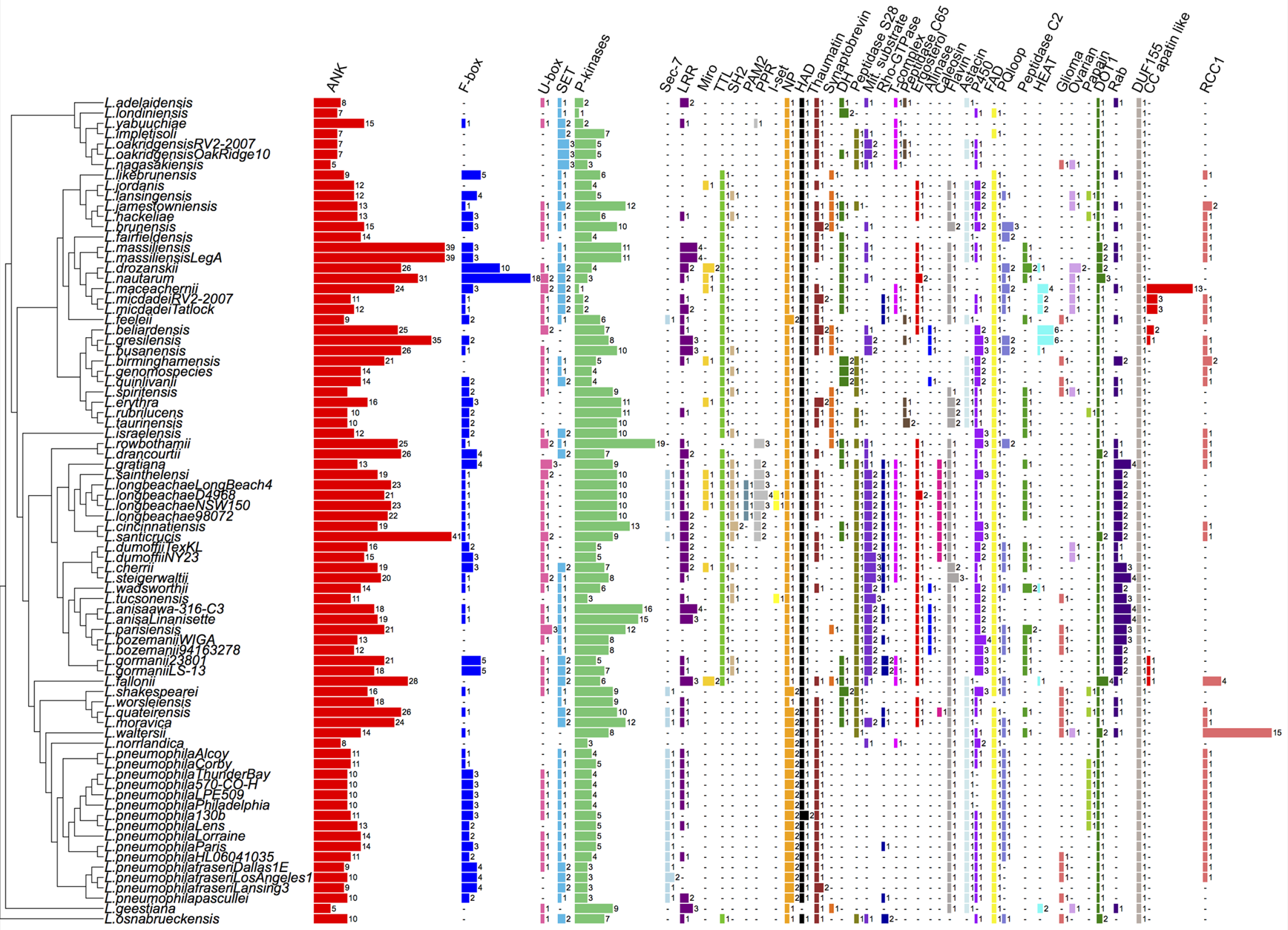

Our analysis revealed the presence of the regulator of chromosome condensation (RCC1) domain in 24 *Legionella* species. Recently it was shown that *L. pneumophila* uses RCC1 repeat containing Dot/Icm substrate, Lpg1976/ Lpp1959 also named LegG1/PieG/MitF to activate Ran and RanBP1, like its eukaryotic homologue, in order to regulate cytoskeletal and mitochondrial dynamics in the host cell (22, 23) Interestingly, *L. waltersii* possessed 14 copies of the Lpg1976 homologue. Phylogenetic analyses of these homologues showed that all 14 *L. waltersii* proteins cluster, indicating that they might have evolved through duplication events (**SI Appendix, Fig. S2**).

One newly identified motif in *Legionella* was the ergosterol reductase ERG4/ERG24 (IPR001171) domain. Ergosterol is the primary sterol in the cell membranes of filamentous fungi, present in membranes of yeast and mitochondria.(24) Importantly, it is also the major sterol of amoebae such as *A. castellanii* and *A. polyphaga,* the natural hosts of *Legionella*(25, 26). We found that 31 *Legionella* species encoded one or two proteins with the ERG4/ERG24 domain (Fig. 2). In fact, the *L. longbeachae* protein (L1o1320) containing this domain showed 56% aa identity to that encoded by the amoeba *Naegleria gruberi* and 30% aa identity to that encoded by *A. castellani* strain Neff. This domain was also present in other amoebae related bacteria such as *Parachlamydia acanthamoebae* and *Protochlamydia naegleriophila*, as well as *Coxiella burnetii.* Phylogenetic analyses suggested that *L. longbeachae* acquired this domain from amoeba (**SI Appendix, Fig. S3A**).

Another newly identified domain is the C-terminal alliinase present in the clades of *L. beliardensis* and *L. anisa* (Fig. 2). The enzyme alliinase is typically present in plants where it is responsible for the production of allicin (27), a defence molecule from garlic (*Allium sativum*). Alliinase can inhibit the proliferation of both bacteria and fungi (28) and thus an effector with this domain that was most likely acquired form plants or other amoeba associated bacteria (**SI Appendix, Fig. S3B**), might help *Legionella* to fight competitor bacteria or fungi in amoebae or in the environment.

An interesting finding was the presence of a Caleosin domain in many strains of the *L. longbeachae* clade (Fig. 2), bacteria that are known to be present in moist soil and dust and thus might co-evolve with plants and fungi. In addition to the *L. longbeachae* clade, this domain is found in only four prokaryotic genomes and in fungi. Caleosin is a lipid-associated protein with a role in the degradation of oil-bodies ubiquitous in plants and fungi (29) and may function in membrane fusion/fission events involved in trafficking between the organelle and storage or transport vesicles. As this domain is found only in Legionella and fungi, the only explanation is *Legionella* acquired it from plant proteins (**SI Appendix, Fig. S3C**). Taken together, our analyses highlight key domains preferentially present in protozoa, fungi, plants or animals that have been acquired by different *Legionella* species.

### A unique case in the prokaryotic world: *Legionella* encode small GTPase-like domains

The Ras-related small GTPase superfamily comprises more than 150 members in humans, which function as key regulators of signal transduction in almost all cellular processes(30). These enzymes bind and hydrolyse GTP to GDP and activate downstream effectors when bound to GTP. The first identified member was the p21-Ras protein, an evolutionary conserved small GTPase that controls cell proliferation, survival and migration through its effector binding at RAF/MAPK and PI3K (31). The Ras protein superfamily is subdivided into at least five distinct branches: Ras, Rho, Rab, Arf and Ran (32). Evolutionarily conserved orthologs are found in *Drosophila, C. elegans, S. cerevisiae, S. pombe, Dictyostelium* and plants (33).

The only Rab-like protein in a prokaryotic genome was reported in the *L. longbeachae* genome sequence (16). However, upon analysis of our 80 *Legionella* strains, we identified 184 small GTPases of which 104 could be classified with a very high confidence as Rab, Ras or Rho like proteins (34 Ras, 71 Rab and one Rho domain) (**SI Appendix, Table S4 and Fig. S4**). Blastp analysis of these proteins in the NCBI database revealed that 149 of the 184 small GTPases of *Legionella* were exclusively present in *Legionella* and eukaryotic organisms (**Table 1**). The Rab domain was localized to different parts of the effector proteins, and a subset of Rab proteins carried additional domains such as U-box domains, ankyrin motifs or F-box domains (Fig. 3A). Alignment of the different Rab domains identified in the *Legionella* genomes revealed that the structural features of eukaryotic Rab domains were conserved among the *Legionella* proteins (**SI Appendix, Fig. S5**).

**Table 1.** Homology of Legionella Rab domains-containing proteins against protozoan Rab proteins

| Domain | Protein | First blast hit | Identity | Coverage | e-value |
| --- | --- | --- | --- | --- | --- |
| Rab | Lade0491 | *Entamoeba histolytica* | 35% | 52% | 4.E-17 |
| Rab | LgoA0634 | *Paramecium tetraurelia* | 33% | 51% | 2.E-19 |
| Rab | Llo3288 | *Ichthyophthirius multifiliis* | 42% | 53% | 4.E-31 |
| Rab | Lstei0814 | *Tetrahymena thermophila* | 34% | 86% | 3.E-26 |
| Rab | Lstei2185 | *Stentor coeruleus* | 38% | 55% | 6.E-29 |
| Rab | Lbir2252 | *Entamoeba invadens* | 32% | 55% | 5.E-15 |
| Rab | Lges1860 | *Entamoeba histolytica* | 34% | 55% | 7.E-25 |
| Rab+ Fbox | Lwad3214 | *Paramecium tetraurelia* | 34% | 35% | 7.E-14 |
| Rab | Lgra2891 | *Guillardia theta* | 36% | 56% | 2.E-19 |
| Rab | Lgra3435 | *Entamoeba histolytica* | 35% | 59% | 2.E-27 |
| Rab | Lma1540 | *Paramecium tetraurelia* | 34% | 55% | 1.E-17 |
| Rab + ank | LmasA3690 | *Oxytricha trifallax* | 34% | 19% | 2.E-19 |
| Rab | Lqua0234 | *Dictyostelium fasciculatum* | 38% | 34% | 1.E-25 |
| Rab | Lquin3026 | *Tetrahymena thermophila* | 34% | 57% | 1.E-19 |
| Rab | Lspi0161 | *Naegleria gruberi* | 34% | 24% | 7.E-24 |
| Rab | Lwal3261 | *Paramecium tetraurelia* | 33% | 85% | 7.E-18 |

Each Rab protein listed in the table represents a different orthologous group. Results are based on blastp searches using the non-redundant NCBI database.

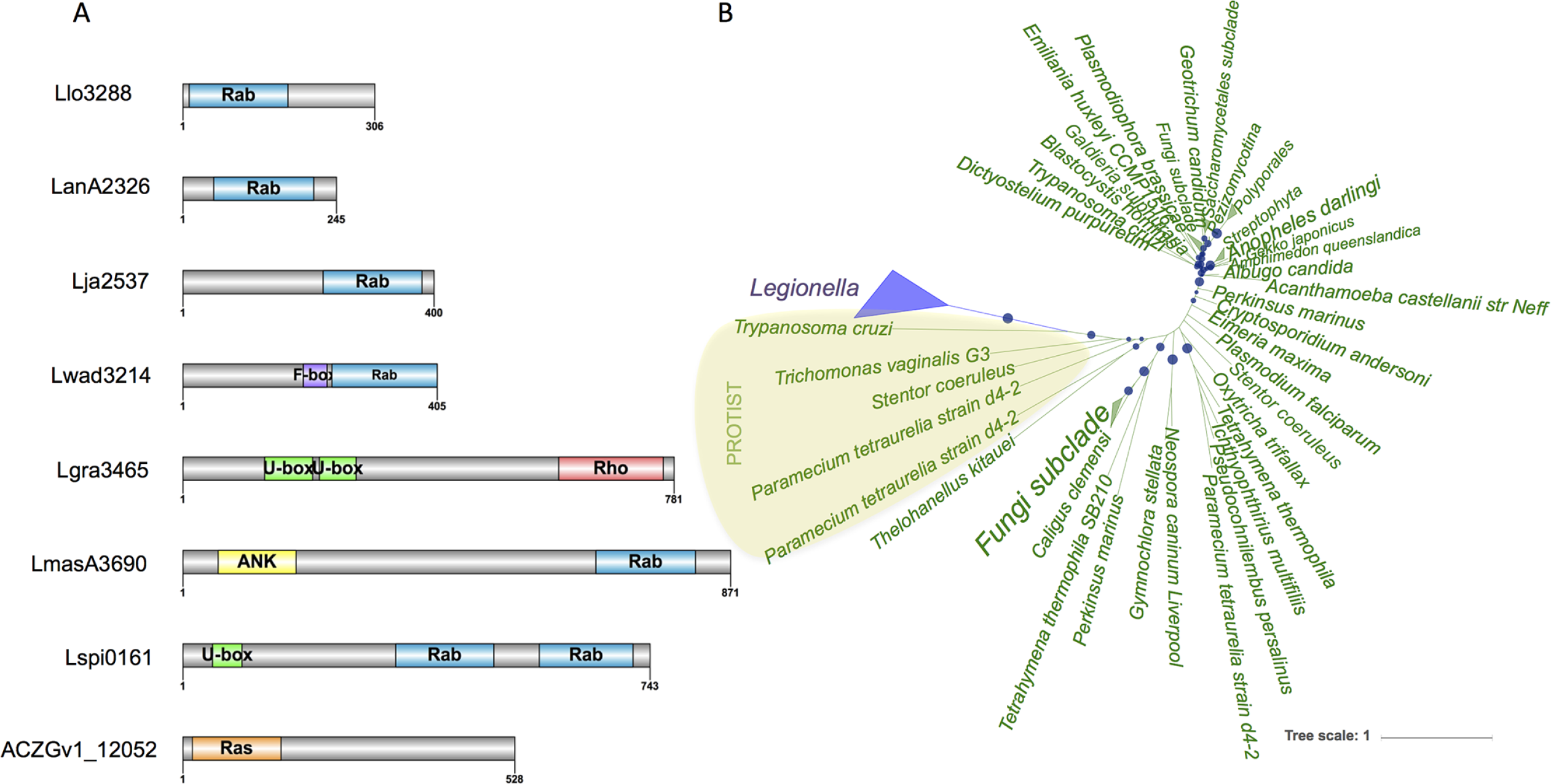

To analyze further the evolutionary history of the Ras-related domains in *Legionella* we undertook phylogenetic analyses of these proteins. For example, the two *L. longbeachae* Rab proteins, Llo1716 and Llo3288, were present in all strains closely related to *L. longbeachae*, suggesting that they and their orthologous share a common origin and evolved from a gene acquired by the ancestor of all these species (**SI Appendix, Fig. S6**). Further phylogenetic analysis of 16 Rab proteins present in eight different *Legionella* species showed that these Rab domains were acquired by HGT, mainly from protozoa (**Fig. 3B and SI Appendix, Fig. S7A-P**). Recently a novel isoform of Rab5D was identified in the *Acanthamoeba polyphaga mimivirus* (APMV) and all group I members of the *Mimiviridae* (34). Phylogenetic analyses suggested that the Rab GTPase was acquired by an ancestor of the *Mimiviridae* family and Rabs from *Mimiviridae, Plasmodium* and few lower eukaryotes form a separate clade (34). Thus, *Legionella* and APMV that both infect the protozoa *Acanthamoeba* encode Rab proteins most likely to mimic and subvert host cell function. To substantiate that these proteins act in the host cell, we determined whether the Rab containing proteins were bona fide substrates of the Dot/Icm T4SS by creating fusion proteins between the 16 different Rab proteins and the catalytic domain of the TEM-1 beta-lactamase (indicated by a star in **SI Appendix, Fig. S6**). Translocation assays were performed using wild type *L. pneumophila* as a surrogate host and compared with an isogenic Dot/Icm mutant (Δ*dotA*). All 16 Rab motif-containing proteins were translocated by *L. pneumophila* but not by the Δ*dotA* mutant (**SI Appendix, Fig. S8**).

### More than 250 different eukaryotic like proteins are encoded in *Legionella* genomes

In addition to modular effectors with eukaryotic domains, the *Legionella* genome encodes proteins that are similar to eukaryotic proteins, many of which are proven effectors of the Dot/Icm T4SS. A wider search for eukaryotic like proteins in the *Legionella* genus identified 2196 eukaryotic like proteins representing more than 400 different orthologous groups that matched better to eukaryotes than to prokaryotes from a total of 6809 different orthologous proteins that matched with eukaryotic proteins. Among these, we identified 156 proteins with a eukaryotic domain, and 210 new eukaryotic-like proteins (**SI Appendix, Table S5**). Furthermore, 152 eukaryotic like proteins detected possess a higher GC content (40%-62%) than the rest of the genome indicating recent HGT. Phylogenetic analysis of selected, newly identified proteins suggested that these were acquired from eukaryotes. As an example, **SI Appendix, Fig. S9** shows the protein LanA0735 from *Legionella anisa*, a species frequently found in artificial water systems. This protein belongs to the pyridine nucleotide-disulfide oxidoreductase family, a subfamily of the FAD dependent oxidoreductase family. LanA0735 showed some similarity to thioredoxin reductase that exists as two major ubiquitous isoenzymes in higher eukaryotic cells, one cytosolic and the other one mitochondrial. The cytosolic form has been implicated in interference with the acidification of the lysosomal compartment in *C. elegans* (35), and thus LanA0735 may help *Legionella* avoid vacuole acidification during infection.

Among the proteins defined as eukaryotic like, two previously described phospholipases of *L. pneumophila*, PlcB (Lpp1411/Lpg1455) and PlcA (Lpp0565/Lpg0502) were identified in our analysis as 90% of the 25 first BlastP hits from eukaryotic proteins. The only other bacteria encoding these two enzymes are *Pseudomonas* and amoebae-associated bacteria. The two enzymes have phospholipase activity (36), but their role in infection is unknown. Here they were predicted as phosphatidylcholine-hydrolyzing phospholipase C. Phosphatidylcholine is a eukaryotic membrane phospholipid that is present in only about 15% of prokaryotic species, in particular bacteria interacting with eukaryotes (37). *L. pneumophila* belongs to the phosphatidylcholine-containing group of bacteria, which includes *Francisella tulurensis* or *Brucella abortus* (38). These pathogens use the phosphatidylcholine synthase pathway exclusively for phosphatidylcholine formation and are thought to depend on choline supplied from the host cell (39). Thus, it is tempting to infer that the role of these enzymes may be to help acquire choline from the host cell.

### Evolutionary history of eukaryotic domains and eukaryotic proteins

It is intriguing that *Legionella* species encode such a diverse repertoire of eukaryotic domains and eukaryotic-like proteins. To understand better this unique feature of the genus we analyzed the evolutionary history of these proteins. After phylogenetic reconstruction of the genus *Legionella* based on the core genome (at least 50% identical) (Fig. 1A), we analyzed the distribution of the eukaryotic motifs and the eukaryotic proteins with respect to the evolution of the genus. For most we found patchy distribution, as the repertoire of these proteins is variable among the different *Legionella* species (Fig. 2). Such a distribution is indicative of gain and loss events during the evolution of the genus. To analyze further how these proteins may have evolved in *Legionella* we selected 25 eukaryotic motifs representing 2,837 different proteins in over 800 orthologous groups and used the program Gloome to analyze the gain and loss events for these proteins. We found that the number of gain events (1,197/69%) considerably exceeded the number of loss events (549/31%), a bias that was even stronger when using parsimony (1,628 gain events *versus* 89 loss events) (**SI Appendix, Fig. S10**). These results were confirmed also when using a more conservative approach by taking a probability cut-off for the stochastic model of 0.8 instead of 0.5, and when analyzing each motif separately.

An exemplary view of this result is shown for four proteins encoding different motifs (U-box and ankyrin repeat, SET domain and ankyrin repeat, astacin domain and allinase domain; Fig. 4). Loss events are indicated by a star and gain events by a dot. The number of gain events exceeds the number of loss events, indicating that in the *Legionella* genus gene acquisition is dominant. Moreover, gene acquisition seems to be an on-going and frequent process in the genus *Legionella* given the high number of events we observed and the fact that most of them are localized in the terminal branches of the tree (**SI Appendix, Fig. S10**). To analyse if eukaryotic-like proteins have the same evolutionary history, we took the sphingosine1-phosphate lyse (*L*pSpl) (40) as an example. Indeed, when running the same analyses this gene also appeared to have been gained multiple times during the evolution of the genus (Fig. 4).

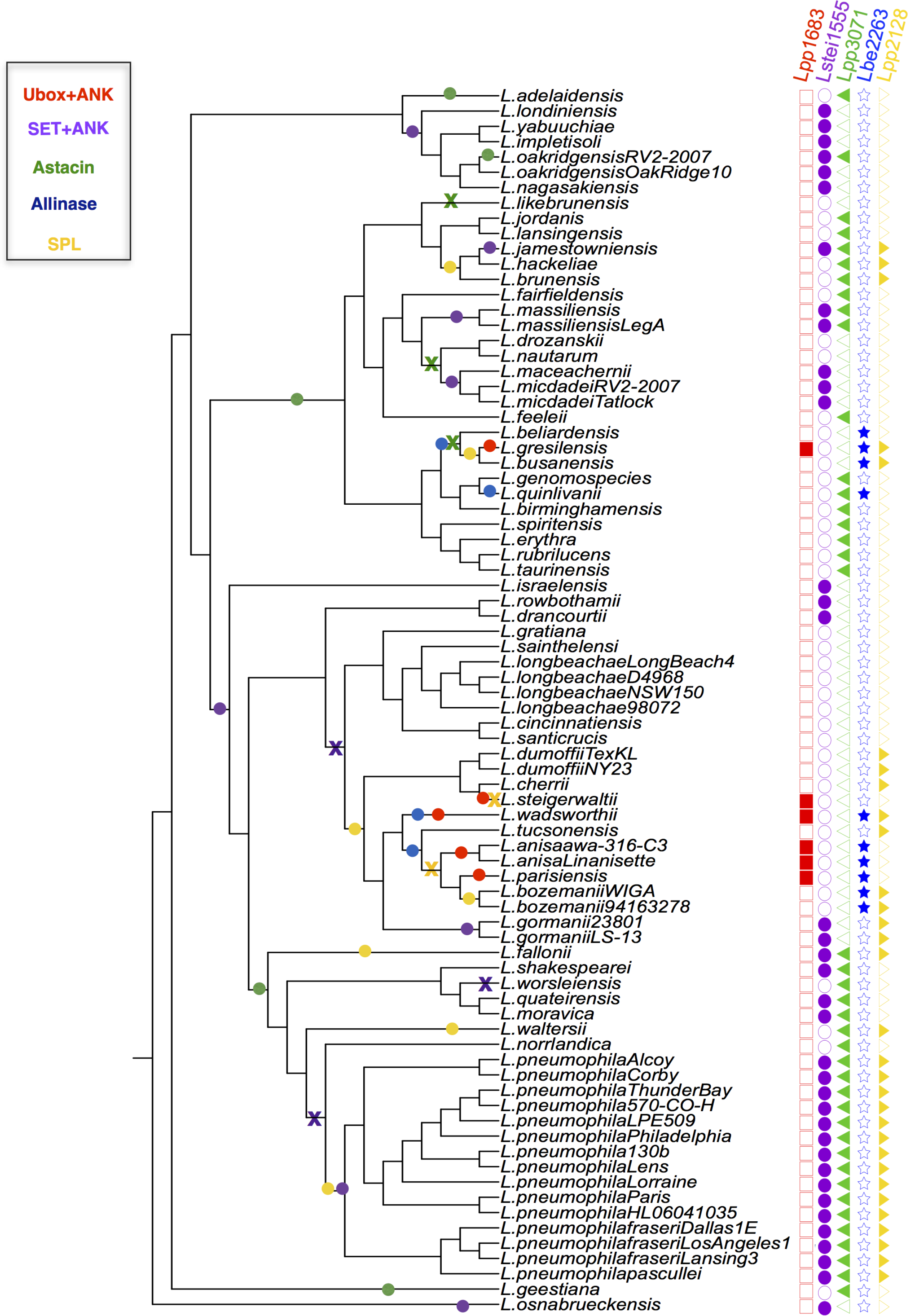

Thus, in comparison to most prokaryotic species analysed to date, more gene gain events are evident than loss events during evolution of the *Legionella* genus, which is also corroborated by the fact that the more ancient genomes (Fig. 1A, cluster 1) are significantly smaller (2.37 - 2.89Mbases).

### The Dot/Icm secretion system is a highly conserved machinery secreting thousands of different proteins

The Dot/Icm T4SS is indispensable for intracellular replication of *L. pneumophila* in both amoeba and macrophages (41). In stark contrast to the high genetic diversity observed in the *Legionella* genomes, the Dot/Icm T4SS is part of the core genome as it is present in all species analyzed and the organization of the constituent proteins is highly conserved, even at the amino acid level. The proteins comprising the secretion machinery show an average amino acid identity of more than 50% and some even more than 90% when compared to the *L. pneumophila* Dot/Icm components (**SI Appendix Fig. S11A and Table S6**). One of the most conserved proteins is DotB, a secretion ATPase (86-100% aa identity) and IcmS, a small acidic cytoplasmic protein (74-98% aa identity). This high conservation is even seen with one of the few non-*Legionella* species that encode a Dot/Icm system, *Coxiella burnetti*.

The only gene of the Dot/Icm system that is not present in all *Legionella* species is *icmR.* IcmR interacts with IcmQ as a chaperone preventing IcmQ self-dimerization (42). Although IcmQ is highly conserved, the gene encoding IcmR is frequently replaced by one or two non-homologous genes encoding for proteins that are called FIR, because they can functionally replace IcmR (43). When overlapping the occurrence of the different FIR genes with the phylogeny of the species, most phylogenetically closely related species share homologous FIR genes (**SI Appendix, Fig. S12**). Apart from two conserved regions (**SI Appendix, Fig. S13**), the absence of sequence homology among FIR proteins indicates that *icmR* is an extremely fast evolving gene and therefore probably under positive selection. The reason why this gene is extremely divergent is still unknown but could be also linked to the high variety of Dot/Icm effectors described in this genus. Thus, except for the FIR genes, the Dot/Icm T4SS is highly conserved and encoded in a very dynamic genetic context.

It has been shown previously, that the more than 300 substrates of the *L. pneumophila* Dot/Icm system are not universally present within the genus *Legionella* as among 38 *Legionella* species only 7 core effectors had been described (14). Surprisingly, although we identified also a core set of 7 substrates these vary when adding the 20 additional genomes sequenced in this study. We confirmed Lpg0103 (VipF), Lpg0107 (RavC), Lpg2300 (LegA3/ankH/ankW), and Lpg2815 (IroT/MavN) as core substrates (**SI Appendix, Fig. S11B and Table S7**). Three of the previously defined core substrates were present as two consecutive genes instead of one, however, this fragmentation might be a sequencing error, and thus these substrates can be considered core substrates (**SI Appendix Table S7**). Contrary to previous reports, we identified three additional core effector genes. These are *lpg1356/lpp1310, lpg2359* and *lpg2936* (**SI Appendix, Fig. S11B and Table S7**). Similarly, to most of the other core substrates, their functions are not known, but each of them encodes a specific motif. Lpg1356 encodes eight eukaryotic Sel-1 motifs similar to LpnE, a *L. pneumophila* virulence determinant that influences vacuolar trafficking (44). It will thus be interesting to elucidate the function of the Lpg1356/Lpp1310 core effector. Lpg2359 and Lpg2936 contain motifs that are related to tRNA metabolism and ribosomal maturation, respectively. Furthermore, 7 other genes are present in all but one, two or four genomes, thus they can similarly be considered as core substrates and might have important functions in host pathogen interactions (**SI Appendix Table S7**).

Interestingly, whereas the number of core Dot/Icm substrates is extremely small, the number and the diversity of predicted Dot/Icm substrates is extremely high. Indeed, through a machine learning approach, Burstein *et al* predicted that the *Legionella* genus would encode 5,885 effectors(14). In contrast, our analyses identified 4,767 proteins with eukaryotic motifs and 2,196 eukaryotic like proteins representing together more than 1,400 different orthologous proteins, and thus more than 1,400 different putative substrates of the Dot/Icm T4SS. If we consider that the orthologues of these proteins in each species are also effectors then the total number of Dot/Icm substrates in the entire genus reaches more than 8000. When adding to the effectors predicted in this study (based on their similarity to eukaryotic domains and proteins), plus the effectors previously described in *L. pneumophila* and their orthologues, the number of effectors rises to almost 16,000 proteins (more than 1,600 orthologous groups) (**SI Appendix, Table S8**). Therefore, the *Legionella* genus has by far the highest number and widest variety of effectors described for an intracellular bacterium. Furthermore, when calculating the growth accumulation curve for Dot/Icm predicted effectors, this number should still increase with the sequencing of new *Legionella* genomes, as the plateau is not reached yet (**SI Appendix, Fig. S11C**).

### The ability to infect human cells has been acquired independently several times during the evolution of the genus *Legionella*

Among the 65 *Legionella* species known, *L. pneumophila* is responsible for over 90% of human disease, followed by *L. longbeachae* (2-7% of cases, except Australia and New Zealand with 30%(45)). Certain *Legionella* species such as *L. micdadei, L. dumoffii* or *L. bozemanii* have once or sporadically been associated with human disease (45), and all other species seem to be environmental bacteria only. The reasons for these differences are not known. To explore whether all species are able to replicate in human cells we chose the human macrophage like cell line THP-1 as model and tested the replication capacity of 47 different *Legionella* species. Infections were carried out in triplicate and colony-forming units were recorded at 24h, 48h and 72h post infection. Levels of intracellular replication were compared to wild type *L. pneumophila* strain Paris and an isogenic non-replicating Δ*dotA* mutant as reference strains (Fig. 5 **and SI Appendix, Fig. S14**). Results were also compared to data previously reported for different *Legionella* species in THP-1, U937 and A549 cells, Mono Mac 6, mouse and guinea pig derived macrophages, or in guinea pigs (**SI Appendix, Table S9**). When results at 72 h after infection were analyzed, 28 of the 47 species tested were impaired for intracellular replication whereas nine species replicated similarly to *L. pneumophila* Paris or better (Fig. 5). These nine species were *L. gormanii, L. jamestowniensis, L. jordanis, L. like brunensis, L. maceachernii, L. micdadei, L. nagasikiensis L. parisiensis,* and *L. tucsonensis*. Interestingly, *L. jamestowniensis,* for which one human case has been reported (46), replicated better than *L. pneumophila* Paris. Indeed, *L. jamestowniensis* productively infects human U937-derived phagocytes. The remaining eight species showed variable replication patterns, being significantly different from *L. pneumophila* Paris only in one or two of the three analyzed time points (**SI Appendix, Fig. S14**). Broadly, the species most frequently reported from human disease (*L. pneumophila L. longbeachae L. micdadei L. bozemanii* and *L. dumoffii*) are also those that replicated robustly in THP-1 cells. The only exception was the *L. dumoffii* strains that were impaired for replication in THP-1 cells but which have been shown to replicate in other cell types and guinea pigs. Taken together, there is a convincing correlation between the frequency of isolation from human disease and the ability to grow in macrophage-like cells.

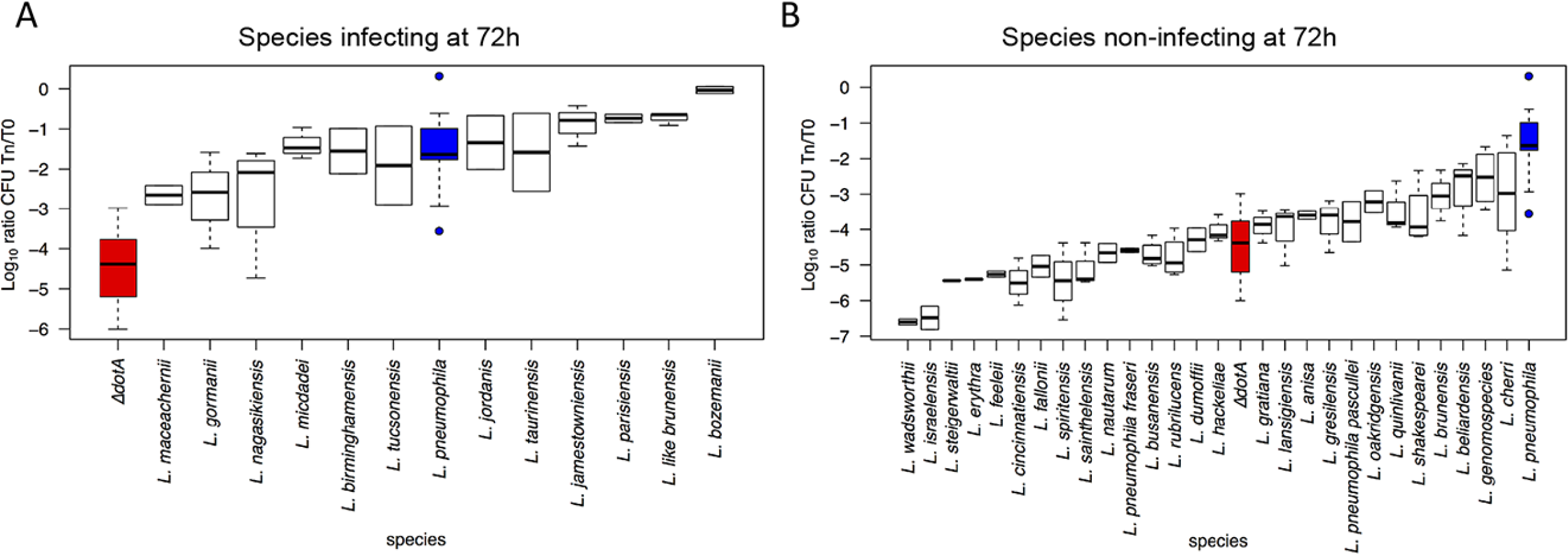

To analyze this further, we overlapped the replication results with the phylogeny of the genus. Apart from the small cluster containing *L. beliardensis*, *L. gresilensis* and *L. busanensis* which were all unable to grow in THP-1 cells, replicating and non-replicating strains were mixed in the phylogeny (**SI Appendix, Fig. S15**). This suggests that the capacity to replicate in human cells has been acquired independently several times during evolution of the *Legionella* genus, possibly as a result of recruiting effectors that allow adaptation to particular niches. To understand whether a specific set of effectors is necessary to infect human cells, we further analyzed the combination of effectors present in the strains isolated from human disease and effectors present in strains capable of replicating in THP-1 cells. Although certain conserved motifs were present, such as ankyrin motifs, F-box or Set domains, no specific set of effectors could be attributed to strains capable of replicating in human cells or isolated from human disease. Thus, the capacity to infect human cells has been acquired independently, several times during the evolution of the genus *Legionella.*

In conclusion, the analysis of 80 *Legionella* strains representing 58 different *Legionella* species has revealed a contrasting picture of the *Legionella* genus. It encodes a highly conserved T4SS predicted to secrete more than 16,000 proteins, of which only 7 are conserved throughout the genus. Together the genomes portray an extremely diverse genus shaped by massive inter-domain horizontal gene transfer, circulating mobile genetic elements and eukaryotic like proteins. Our in-depth analyses of eukaryotic features of the *Legionella* genomes identified 137 different eukaryotic domains of which Rab or Ras domain-containing proteins were quasi unique to the genus *Legionella.* The secretion assays undertaken for 16 of these Rab or Ras domain-containing proteins confirmed that these were translocated Dot/Icm effectors. In addition to the eukaryotic domains, we identified 210 orthologous groups of eukaryotic like proteins. If all these proteins in the different species and their orthologues are taken into account, we found more than 8,000 proteins that have been shaped by inter-domain horizontal gene transfer in the genus *Legionella.* Thus, to our knowledge the genus *Legionella* contains the widest variety and highest number of eukaryotic proteins and domains of any prokaryotic genus genome analyzed to date. Analyzing more strains per species will probably discover new unknown effectors increasing our knowledge of the set of tools used by *Legionella* to infect eukaryotic cells. Although eukaryotic proteins and domains were a universal feature of the genus *Legionella*, the repertoire of these proteins for each species was different. Surprisingly, even when the same motif was present in different species, these were often present in different proteins with no orthology. In accordance with this finding, our evolutionary analysis of the presence/absence of these domains and proteins suggested that these proteins were mostly acquired through gene gain events.

When exploring the replication capacity of 47 different *Legionella* species in human macrophage-like cell line THP-1, we found that the 23 species were capable of replicating in THP-1 cells. However, these did not cluster in the phylogeny, indicating that the capacity to replicate in macrophages can be achieved by different combinations of effectors, and this capacity has been acquired several times during the evolution of the *Legionella* genus. As humans are an accidental host for *Legionella*, the capacity to replicate in macrophages may also have been obtained by a coincidental acquisition of different virulence properties initially needed to adapt to a specific natural host, such as amoebae. Indeed, due to the high conservation of key signaling pathways in professional phagocytes such as amoebae and human macrophages, different combinations of effectors may allow *Legionella* species to infect higher eukaryotic cells by chance.

Here we show that all *Legionella* species have acquired eukaryotic proteins that likely modulate specific host functions to allow intracellular survival and replication in eukaryotic host cells. At a certain point, the evolution of a combination of effector proteins that allow replication in human cells may inadvertently lead to the emergence of new human pathogens from environmental bacteria.

## Material and Methods

The materials and methods are described at length in *SI Appendix.* This includes: Sequencing and assembly, sequence processing and annotation, pan/core genome, ortholog and singleton definition, phylogenetic reconstruction and evolutionary analysis, phylogenetic analyses of Rab and eukaryotic-like proteins, infection assays, statistical analysis, and translocation assays. The raw sequence reads were deposited in the European Nucleotide Archive (study accession number PRJEB24896).

## Acknowledgements

We would like to thank Tim P. Stinear for critical reading of the manuscript and helpful comments and we acknowledge the receipt of 53 different *Legionella* strains from the Collection of the Institut Pasteur (CIP). Work in the CB laboratory is financed by the Institut Pasteur, the grant n°ANR-10-LABX-62-IBEID and the Fondation pour la Recherche Médicale (FRM) grant N° DEQ20120323697.

## Author contributions

SJ and LGV, ELH contributed to sample collection and strain analyses DNA extraction and sequencing; CR, DC, SM, AEPC, MR, SP, SR, JD, JC, SDD, GNS to functional experiments, data analyses and interpretation. The manuscript was written by LGV and CB with input from co-authors. The project was conceived, planned and supervised by LGV, GD, GF and CB.

